# Smartphone-assisted real-time estimation of chlorophyll and carotenoid contents in spinach following the inversion of red and green color features

**DOI:** 10.1101/2021.03.06.434237

**Authors:** Avinash Agarwal, Piyush Kumar Dongre, Snehasish Dutta Gupta

**Affiliations:** Agricultural and Food Engineering Department, Indian Institute of Technology Kharagpur, Kharagpur - 721302, India; Rajendra Mishra School of Engineering Entrepreneurship, Indian Institute of Technology Kharagpur, Kharagpur - 721302, India

**Keywords:** Image analysis, Chlorophyll, Carotenoid, RGB color model, Smartphone

## Abstract

**Purpose:** Chlorophyll (Chl) content is a reliable indicator of leaf nitrogen content and plant health status. Currently available methods for image-based Chl estimation require complex mathematical derivations and high-throughput imaging set-up along with multiplex image-preprocessing steps. Further, the influence of carotenoid (CAR) content has been largely ignored in the process. The present study describes a smartphone-based leaf image analysis method for real-time estimation of Chl content and Chl/CAR ratio.

**Methods:** Color features were obtained from RGB (red, green, blue) images of spinach leaves using a smartphone, and inverse R and G values were calculated. Correlation analysis of color indices and photosynthetic pigment (PP) contents was performed, followed by principal component analysis (PCA). Linear mathematical modeling was performed for describing regression equations for predicting PP contents.

**Results:** 1/R and 1/G showed strong positive linear correlation (0.93 < *r*^*2*^ < 0.96) with Chl and CAR contents, respectively. Furthermore, 1/R+1/G and [1/R]/[1/G] presented strong positive linear correlation with Chl + CAR (*r*^*2*^ = 0.95) and Chl/CAR (*r*^*2*^ = 0.88), respectively. PCA confirmed the association of color indices with the respective PP features, which were subsequently estimated using the correlation models. A smartphone-based companion application was developed using the linear models for non-invasive, real-time estimation of Chl content and Chl/CAR ratio.

**Conclusion:** The ratios 1/R and 1/G indicate the contents of Chl and CAR via linear models. The smartphone application developed using the linear models enables real-time estimation of Chl content and Chl/CAR ratio without complicated image preprocessing steps or mathematical derivations.

## 1. Introduction

Chlorophyll (Chl) content is considered a reliable indicator of plant health status due to its central role in photosynthesis, as well as its close association with leaf nitrogen content (Kawashima and Nakatani 1998; Carter and Knapp 2001; Rorie et al. 2011 ; Wang et al. 2019). Although the use of Chl meters has been adopted for rapid non-destructive assessment of Chl content, the high cost of such devices has limited their usage significantly. Since Chls are directly associated with leaf coloration, analysis of leaf digital color features is becoming a popular cost-effective tool for non-destructive, real-time assessment of Chl content (Kawashima and Nakatani 1998; Pagola et al. 2009; Rorie et al. 2011; Agarwal and Dutta Gupta 2018). A variety of color indices derived from leaf digital images have been examined with the aim of finding the most suitable indicators of Chl content. Use of red, green, and blue (RGB) color components has been reported most frequently for this purpose (Kawashima and Nakatani 1998; Pagola et al. 2009; Yadav et al. 2010; Vollmann et al. 2011; Dutta Gupta et al. 2013; Riccardi et al. 2014; Rigon et al. 2016; Dutta Gupta and Pattanayak 2017; Agarwal and Dutta Gupta 2018; Hassanijalilian et al. 2020).

Studies with RGB features have unequivocally reported a negative relation of Chl content with R and/or G color indices. Notably, data plots presented in various reports suggest a logarithmic decline in both these color features with increasing Chl content (Yadav et al. 2010; Hu et al. 2013; Riccardi et al. 2014; Rigon et al. 2016), indicating lower sensitivity of the R and G models at higher pigment contents. Other color indices such as R-B, G-B (Kawashima and Nakatani 1998), R+B, R+G, B+G, R+G+B (Hu et al. 2013), R/B, and G/B (Baresel et al. 2017) have also been shown to correlate well with Chl content. However, theoretical explanations for all such correlations from the physiological and photochemistry perspectives may not be possible. Investigations employing machine learning for image-based prediction of Chl content (Dutta Gupta et al. 2013; Vesali et al. 2015, 2017; Dutta Gupta and Pattanayak 2017; Odabas et al. 2015, 2017; Mohan and Dutta Gupta 2019; Hassanijalilian et al. 2020) inherently employ black-box algorithms, and hence, their exact method of computation may not be easily defined. Furthermore, various methods have been adopted to compensate for the interference by ambient light conditions during leaf image acquisition (Hu et al. 2010; Rorie et al. 2011; Riccardi et al. 2014; Confalonieri et al. 2015; Rigon et al. 2016; Mohan and Dutta Gupta 2019). However, such methods either possess limited practical feasibility or make data analysis more complicated, creating the need for a more practicable process.

Although digital image analysis techniques for predicting Chl content aim to interpret changes occurring in leaf coloration, only a few studies (Hu et al. 2013; Rigon et al. 2016) have addressed the contribution of carotenoid (CAR) content. It has been well established that Chls degrade more rapidly than CARs in stressed and senescent leaves (Amir-Shapira et al. 1987; Biswal 1995; Bertrand and Schoefs 1999), causing a change in the relative abundance of Chls and CARs, the factor responsible for the distinct change in leaf color from green to yellow. Thus, the combined impact of Chl and CAR contents on leaf color necessitates their simultaneous assessment for accurate analysis of leaf digital color features.

The present study describes a simple and cost-effective smartphone-based leaf image analysis technique for the simultaneous estimation of Chl and CAR contents, without the need of any specialized equipment, complex mathematical analysis, or image modification. A simple and reproducible fixed light input photography (FLIP) technique was adopted for capturing leaf images with a smartphone, ensuring no interference from ambient light conditions. Comparative analysis of Chl and CAR contents with the inverse of R and G color features was carried out to evaluate their potential as predictors of photosynthetic pigment (PP) contents. Based on the close-fit regression models, a smartphone application was developed for estimating PP content and assessing plant health status.

## 2. Materials and methods

### 2.1. Plant material and growth conditions

Spinach (*Spinacia oleracea* L. cv. “All Greens”) seeds were purchased from Sutton and Sons Pvt. Ltd. (Kolkata, India). Seeds were soaked overnight in sterile distilled water and subsequently suspended in a cool, dark spot for 48 h using a moist muslin cloth. Germinated seeds were sown in three plastic trays containing Soilrite-mix (perlite + vermiculite + sphagnum peat moss 1:1:1) purchased from Keltech Energies Ltd. (Bangalore, India). Trays were kept in a plant growth chamber under cool-white fluorescent lamps with a photoperiod of 16 h, for 10 days. Subsequently, each tray was subjected to a different photoperiod, viz. 0, 6, and 16 h, for the next 10 days. Light deprivation was carried out to induce degreening via dark-induced senescence, as reported previously (Keech et al. 2007). Trays were irrigated with sterile distilled water intermittently. A total of fifty upright leaves having pale yellow (senescent) to dark green (healthy) coloration were visually selected from the 20-day old seedlings, and serially labelled before image acquisition.

### 2.2. Leaf image acquisition by FLIP method

The FLIP method adopted for acquiring leaf images using a smartphone utilized the camera flash as the light source in conjunction with a customized smartphone-mountable enclosed image acquisition chamber (Fig. 1). This setup allows image capture under uniform lighting, irrespective of the external light environment, and may be used indoors as well as in the field.

**Fig 1.**
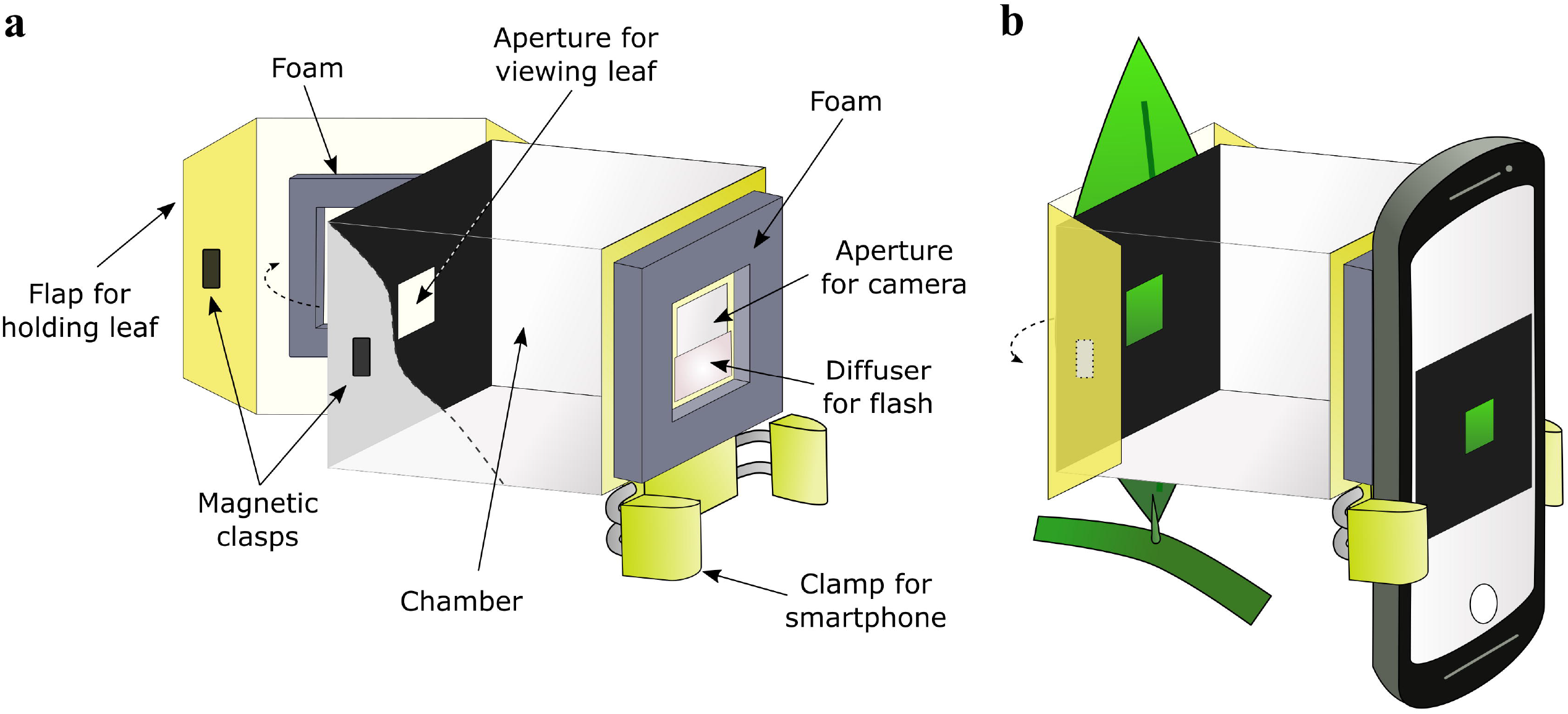
Schematic representation of the structure (a) and mode of operation (b) of the fixed light input photography (FLIP) method adopted for acquiring leaf images via a smartphone. The FLIP chamber and the flap for holding the leaf sample were made with an opaque white, flexible plastic sheet (thickness ∼0.3 mm). Length of the chamber (12 cm) was kept greater than the minimum distance of focusing for the camera to ensure sharp images. Opposite ends along the length of the chamber had openings (2 × 2 cm2) for the camera and for viewing the leaf sample, respectively. The lower half of the opening for the camera was fixed with a white translucent sheet which acted as a diffuser for the camera flash. The opening on the opposite side of the chamber was covered with a matte black paper having a 1 cm2 aperture for viewing the leaf. Dark grey foam (thickness ∼2 mm) was used for bordering both the openings to prevent external light from interfering during image acquisition. The outer surface of the chamber was covered with black paper, and the inner surface with white paper to ensure zero interference from external light. The chamber was attached to a commercially available smartphone holder (clamp) as depicted

Images were captured using a Moto G Turbo Edition android smartphone (Motorola Inc., USA), with camera specifications: 13 megapixels, *f*/2.0 aperture, 76° field of view, dual LED flash. Photographs were taken using default settings, without selecting any image enhancement options. Leaves were photographed without being detached from the plant (Fig. 1b).

### 2.3. Estimation of Chl and CAR contents

Leaves were immediately excised and homogenized in ice-cold 80% (v/v) acetone. Homogenates were centrifuged at 5000 rpm at 4 °C for 15 min. The supernatants were used for spectrophotometric determination of total Chl and total CAR contents as described by Lichtenthaler (1987).

### 2.4. Extraction of color features

Leaf images were transferred to a computer. Mean R, G, and B values were obtained from the region of image containing the leaf (250 × 250 pixels) using ImageJ software (https://imagej.nih.gov/ij/). R and G values were used to obtain the inverse color indices 1/R and 1/G, as well as their ratio [1/R]/[1/G] and sum 1/R+1/G.

### 2.5. Statistical analysis

The relation of color indices with different PP features, viz., total Chls, total CARs, total PP content (Chl + CAR), and Chl/CAR ratio, was analyzed using R software (ver. 4.0.3) (https://www.r-project.org/) in the R Studio (ver. 1.3.1093) environment (https://rstudio.com/). Pearson’s correlation coefficient (*r*) between the color indices and PP features was calculated using the “*cor*” function. The “*lm*” function was used for obtaining linear regression models, with color indices as the independent variable (predictor) and PP features as the target (unknown) variable. Level of significance, coefficient of determination (*r*^*2*^), and root mean squared error (RMSE) values of the models were obtained by using the “*summary*” command. Variation in sensitivity of the prediction models with the magnitude of predictor was assessed by performing first order differentiation using the “*D*” function. The lowest and highest observed values of the predictor were substituted in the differential equations to obtain the initial and final rates of change, respectively. Principal component analysis (PCA) was performed using the “*prcomp*” function to assess the efficacy of the selected color indices, viz., 1/R and [1/R]/[1/G], in segregating the healthy and senescent leaves. PCA biplots were generated using the “*ggbiplot*” package. A similar PCA biplot was prepared for segregating leaf samples based on Ch l content and Chl/CAR values for comparison.

### 2.6. Development of smartphone application

A native smartphone application, titled “Plant Health Assessment”, was developed for leaf image acquisition, extraction of RGB features from the region of interest, and estimation of Chl content and Chl/CAR ratio using the linear prediction models. Based on the close relation between Chl content and plant health status, an additional parameter called the “Plant Health Score” (PHS) was incorporated into the application, which was calculated as follows:

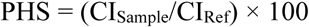

where CI indicates the color index showing a strong correlation with Chl content. CI_Sample_ and CI_Ref_ indicate the color index values for the sample leaf and the reference, respectively. PHS assigns a hypothetical score (out of hundred) to the target sample by comparing its color features with a reference “healthy” leaf selected by the user from the options provided in the application.

The application was developed using Java language and Android Software Development Kit in the Android Studio. A third-party library called Glide (https://github.com/bumptech/glide) was used for selecting the region of interest. The algorithm for the application has been presented in Supplementary Fig. S1. All computations and analyses of the images during the implementation were done in Java, without any external library. The size of the software installation package is around 11 MB. The application is compatible with any android device having Android Jelly Bean (API level 16) or above. The application is in compliance with all the guidelines of Google Play Store, and has been uploaded for testing.

The application was tested by estimating the Chl content, Chl/CAR ratio, and the PHS of the spinach leaf images analyzed previously.

## 3. Results

### 3.1. Comparison of color indices with PP contents

Chl content showed a strong positive linear correlation (*r*^*2*^ = 0.938) with 1/R (Fig. 2a). Similarly, CAR content had a strong positive linear relation (*r*^*2*^ = 0.951) with 1/G values (Fig. 2b). Further, total photosynthetic pigment content, i.e., Chl + CAR, and the ratio of Chl and CAR were strongly correlated with 1/R+1/G and [1/R]/[1/G] values, respectively (Fig. 2c, d). This corroborates the relation between the PP contents and the respective color indices, and implies that 1/R and 1/G accurately represent the abundance of Chls and CARs in leaves. Other comparisons between Chl and CAR contents, and the different color indices have been presented in Supplementary Fig. S2.

**Fig 2.**
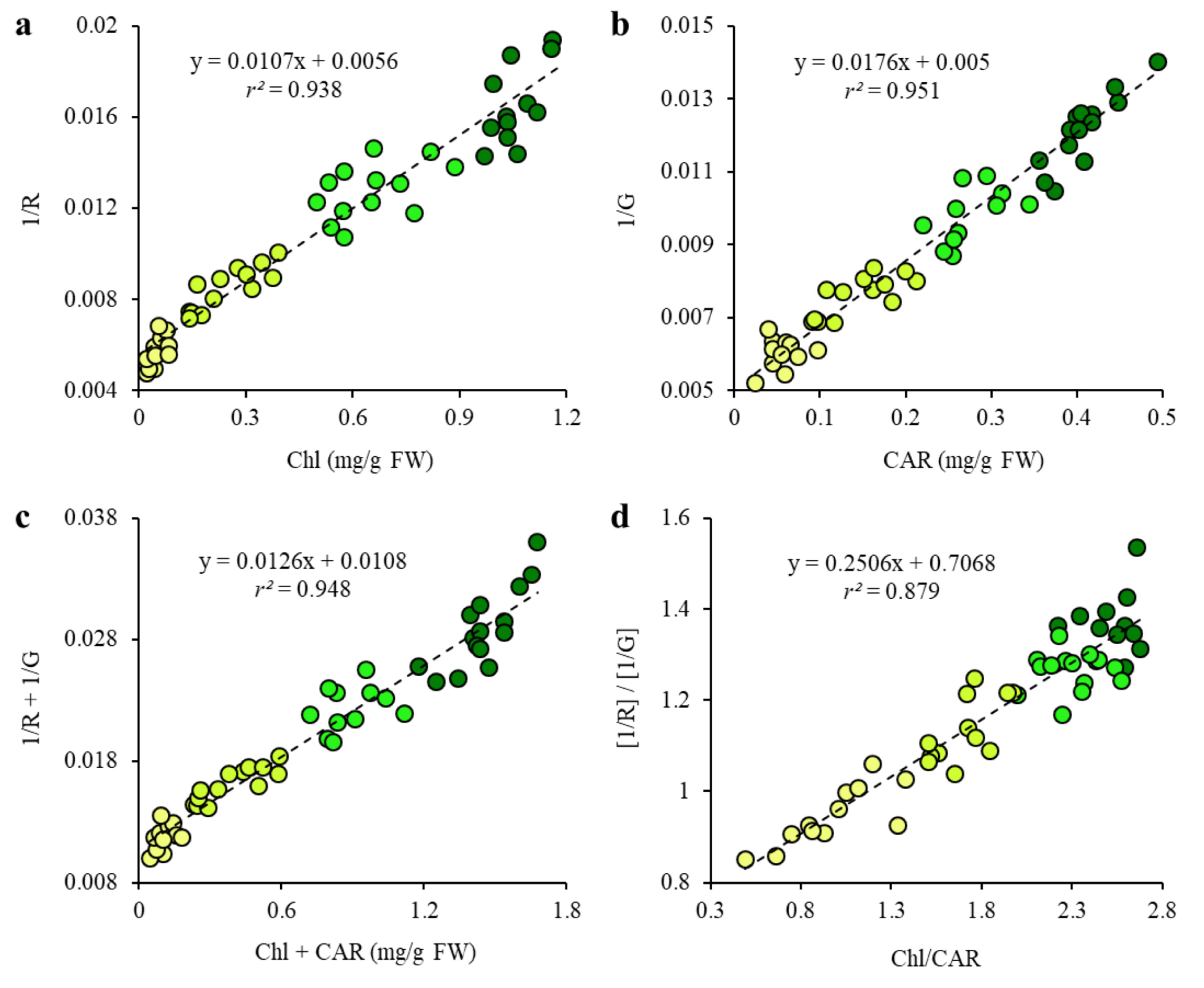
Relation of photosynthetic pigment contents with the red (R) and green (G) digital color features. Correlation between chlorophyll content (Chl) and 1/R (a), carotenoid content (CAR) and 1/G (b), total photosynthetic pigment content (Chl + CAR) and 1/R+1/ G (c), and Chl/CAR ratio and [1/R]/[1/G] (d) (*n* = 50, *p* < 0.001). Shades from light to dark represent lower to higher PP contents

The PCA biplot obtained using the total Chl content and Chl/CAR showed a clear segregation of healthy and senescent leaves (Fig. 3a). A markedly similar clustering of leaf samples was observed in the PCA biplot generated using 1/R and [1/R]/[1/G] color features (Fig. 3b), indicating that these color indices can be used simultaneously to reliably distinguish between the healthy and stressed plants.

**Fig 3.**
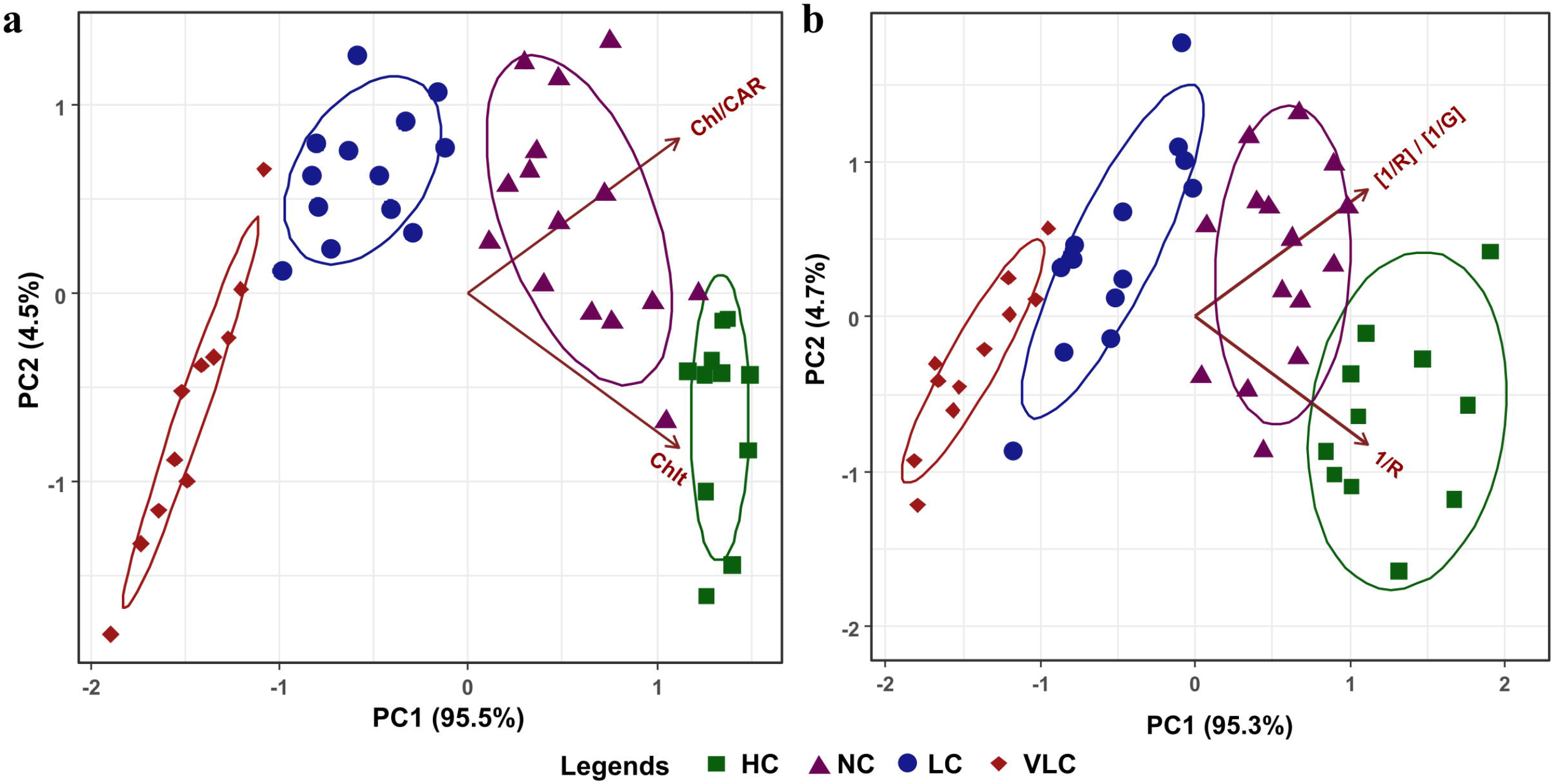
Principal component analysis (PCA) biplots obtained using photosynthetic pigment contents (a) and color features (b). Arrows indicate the variables; Chl, chlorophyll content; Chl/CAR, ratio of chlorophylls and carotenoids; 1/R, inverse of red color feature; 1/G, inverse of green color feature. HC, NC, LC, and VLC indicate leaf samples with high (>1 mg/g FW), normal (>0.5 mg/g FW, <1 mg/g FW), low (>0.1 mg/g FW, <0.5 mg/g FW), and very low (<0.1 mg/g FW) chlorophyll contents, respectively. PC1 and PC2 indicate the first two principal components. Values within parentheses beside each PC indicate the percentage of variance explained by the corresponding PC

### 3.2. Prediction of PP contents using color indices

Model equations for predicting Chl content using R and 1/R, CAR content using G and 1/G, Chl + CAR using 1/R+1/G, and Chl/CAR using [1/R]/[1/G] have been presented along with the *r*^*2*^, RMSE, and the initial and final rates of change (Table 1). Exponential equations used for predicting Chl and CAR contents had comparable RMSE values as the linear equations, indicating similar accuracy of prediction. However, in contrast to the linear models, the initial and final rates of change of the exponential models for predicting Chl and CAR contents showed considerable variation.

**Table 1.** Equations for estimation of photosynthetic pigment (PP) features, i.e. chlorophyll (Chl), carotenoid (CAR), and Chl+CAR contents, and Chl/CAR ratio using the color indices obtained from leaf digital images, coefficients of determination (*r*^*2*^), root mean square errors (RMSE), and the initial and final rates of change in the color index with respect to the PP content.*

| PP feature<br>(y) | Colour<br>index (x) | Equation | $r^2$ | RMSE | Initial rate<br>of change<br>($dy/dx_i$) | Final rate<br>of change<br>($dy/dx_f$) |
| --- | --- | --- | --- | --- | --- | --- |
| Chl | R | $y = 4.825\exp^{(-0.0257x)}$ | 0.958 | 0.108 | -0.0359 | $-6.86 \times 10^{-4}$ |
| | 1/R | $y = 87.48x - 0.454$ | 0.938 | 0.110 | 87.48 | 87.48 |
| CAR | G | $y = 2.814\exp^{(-0.0233x)}$ | 0.925 | 0.037 | -0.0209 | $-8.17 \times 10^{-4}$ |
| | 1/G | $y = 53.97x - 0.26$ | 0.951 | 0.031 | 53.97 | 53.97 |
| Chl+CAR | 1/R + 1/G | $y = 75.5x - 0.778$ | 0.948 | 0.122 | 75.5 | 75.5 |
| Chl/CAR | [1/R]/[1/G] | $y = 3.51x - 2.25$ | 0.879 | 0.216 | 3.51 | 3.51 |

\* $n = 50$ , $p < 0.001$ ; $\exp = 2.7183$ (approximately).

### 3.3. Smartphone application for plant health assessment

The smartphone application enabled the measurement of RGB values as well as estimation of Chl content and Chl/CAR ratio directly from leaf images (Fig. 4a, b). The linear regression model with 1/R (Table 1) was used for estimating Chl content using the application, whereas Chl/CAR ratio was estimated using the linear model with [1/R]/[1/G] (Table 1). The application was also designed to provide qualitative indication regarding Chl/CAR ratio as very low, low, medium, high, or very high (Fig. 4a). Screenshots of the application user-interface have been presented in Supplementary Fig. S 3.

**Fig 4.**
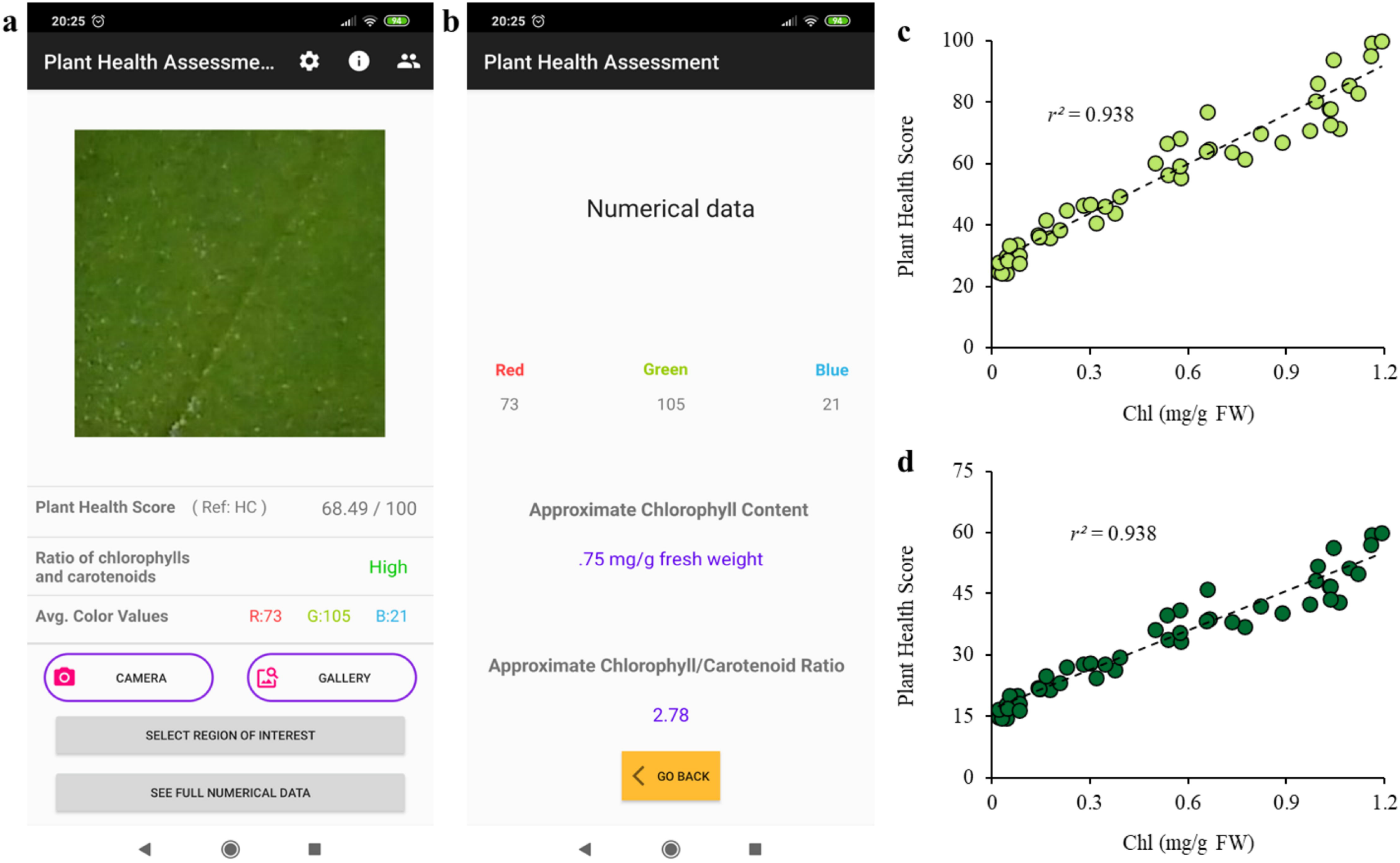
Screenshots of the “Plant Health Assessment” smartphone application user interface (a, b), and the plant health scores calculated from leaf images using the smartphone application with “high chlorophyll” (HC) (c) and “very high chlorophyll” (VHC) (d) as the references (n = 50, *p* < 0.001). Chl, chlorophyll content

The color index 1/R was selected for calculating the PHS in the application as this color index showed strong positive linear correlation (Fig. 2a) and possesses plausible theoretical association with Chl content (discussed later). PHS was calculated using the experimental spinach leaf images by selecting the “high Chl” (HC; R = 50; 1/R = 0.02) and “very high Chl” (VHC; R = 30; 1/R = 0.033) options as the reference in two independent runs (Fig. 4c, d). The VHC reference assigned lower PHS, whereas HC reference gave higher scores to the same leaves. Both curves showed the same correlation with Chl as they were calculated using the same linear equation.

## 4. Discussion

RGB color features of digital images represent the reflection of light in the red, green, and blue wavebands by an object (Palus 1998). Intensity of reflection from all three wavebands collectively determines the color of the object. Since PPs are capable of selective absorption among these three wavebands, RGB color features have the potential to provide reliable information regarding PPs present in leaves.

In the present study, Chl and CAR contents showed a strong linear correlation (Supplementary Fig. S2a), whereas Chl/CAR exhibited an asymptotic increase with respect to Chl + CAR content (Supplementary Fig. S2b). Earlier studies have reported that yellowing or senescing leaves undergo faster decline in Chl content as compared to CAR, termed as CAR retention (Merzlyak et al. 1999, 2003), which leads to low Chl/ CAR values at low Chl contents, as observed in the present study. Spectral reflectance analyses by Merzlyak et al. (1999, 2003) and Gitelson et al. (2002) indicated that healthy leaves have low total reflectance, and reflect green photons more than red photons, whereas senescing leaves have high total reflectance, and reflect photons in the red and green wavebands almost equally. Merzlyak et al. (2003) reported changes in reflectance at 500 nm (green light, *R*_500_) and 678 nm (red light, *R*_678_) in healthy and senescent leaves. They observed that leaf samples with high Chl content had higher *R*_500_ values than *R*_678_ values. Though senescence caused simultaneous increase in both *R*_500_ and *R*_678_ values, the *R*_678_ value increased at a faster rate than *R*_500_, causing the two values to become almost equal in yellow (senescent) leaves. In the present study, a similar trend was noted in the R and G values; the decrease in PP contents resulted in an increment in R and G values (Supplementary Fig. S2f, g), along with a concomitant increase in the R/G ratio, with values reaching 0.9 and 1.2 in the senescent leaves (Supplementary Fig. S2h). High degree of similarity between the *R*_678_ and *R*_500_ reflectance features reported earlier and the R and G color features of the present study indicates strong analogy between the leaf reflectance parameters and the respective digital color features. Chlorophyll reflectance studies by Sims and Gamon (2002) and Gitelson et al. (2003) also portrayed the asymptotic reduction in red light reflectance with increasing Chl contents, which is comparable to the findings of the present study (Supplementary Fig. S2f).

Since R and G values represent the amount of red and green light reflected by the leaves, respectively, the inverse values of R and G color features, i.e., 1/R and 1/G, may be considered analogous to light absorbance in the corresponding wavebands. Analyzing 1/R and 1/G yielded a strong positive linear relation with Chl and CAR contents (Fig. 2a, b), clearly portraying the characteristic absorption of light in the red and green wavebands by the respective PPs. This representation significantly simplifies the relation between leaf color features and PP contents, and provides a straightforward method for estimating the abundance of PPs in leaf samples. Further, the sum of 1/R and 1/G also showed good correlation with Chl + CAR content (Fig. 2c), supporting the previous observation.

The analogy of 1/R and 1/G with light absorption by Chls and CARs in the red and green wavebands, respectively, is further substantiated by the strong positive correlation between Chl/CAR ratio and [1/R]/[1/G] (Fig. 2d). In previous studies, estimation of Chl/CAR ratio was carried out via comparison of reflectance in Chl and CAR absorbance regions (Merzlyak et al. 1999, 2003; Sims and Gamon 2002). Similarly, in a recent study by Gitelson (2020), ratio of Chl and CAR was estimated by comparative analysis of absorption coefficients at 500 and 660 nm using leaf reflectance data collected from seven plant species. Use of [1/R]/[1/G] takes an analogous, but simpler approach for estimating Chl/CAR. Furthermore, simultaneously using 1/R and [1/R]/[1/G] allows effective segregation of healthy and senescent leaves, and shows a similar distribution of samples as that observed in the PCA biplot generated using Chl and Chl/CAR (Fig. 3). This suggests that the inverse color indices, i.e., 1/R and 1/G, have the capacity to accurately represent the abundance of PPs, and may be used for distinguishing healthy and stressed/senescent leaves via machine vision.

Testing the smartphone application yielded similar RGB values, Chl content, and Chl/CAR ratio (data not shown) as that observed during the computer-based analyses, indicating that the application was working as expected. Slight deviations were observed due to minor variations in RGB values (< 5%) recorded by the application. PHS obtained using the two references (Fig. 4c, d) showed strong correlation with Chl content (*r*^*2*^ = 0.938). Since Chl content is considered a primary indicator of plant health, the PHS parameter defined in this study may be adopted as general indicator of relative plant health status, similar to chlorophyll meter readings. Vesali et al. (2015) developed a smartphone application wherein information from multiple color spaces obtained from images of corn leaves was used for predicting Chl content in terms of Chl meter readings via a linear model or an artificial neural network. An application developed by Tao et al. (2020) directly compared the color features of the target leaf with sample leaves in real time for estimating nitrogen content. The application developed in the present study adopts a distinct, but simpler approach for analyzing leaf color information for estimating Chl and CAR contents, giving an insight into the plant’s health status.

## 5. Conclusion

Owing to its close association with the absorption and reflectance patterns of different PPs, leaf color information present in R and G values of the RGB color space provides information regarding Chl and CAR contents with high fidelity. Using 1/R and 1/G values derived from leaf images allows reliable estimation of total and relative contents of PPs over a wide range, without the need of complicated mathematical transformations or image manipulations. The smartphone application developed as a part of this study further simplifies the process by directly providing the RGB values, as well as Chl content, Chl/CAR ratio, and PHS. The present study provides a preliminary insight into the potential of using the reciprocals of leaf RGB color features for easy estimation of plant health status. As the present study was carried out using only one plant species and one model of smartphone, future investigations exploring the application of this technique using different plant species and other smartphones or digital cameras shall further establish its practical feasibility. Thus, further studies are warranted to confirm the relationships as derived from 1/R and 1/G for their reliability as universal indicators of Chl and CAR contents.

## Declarations

### Ethics approval

Not applicable. **Consent to participate:** Not applicable. **Consent for publication:** Not applicable.

### Availability of data and material

Available from the corresponding author upon reasonable request.

### Code availability

Statistical analysis was performed using standard commands in R. The application developed in this study is freely available in the Google App store.

## Acknowledgements

None.

## Authors’ contributions

Avinash Agarwal: Conceptualization, Methodology, Formal analysis, Investigation, Writing-Original draft preparation, Writing-Reviewing and Editing, Visualization.

Piyush Kumar Dongre: Software.

Snehasish Dutta Gupta: Supervision, Conceptualization, Writing-Reviewing and Editing.

