## Supplemental figures for "Smartphone-assisted real-time estimation of chlorophyll and carotenoid contents in spinach following the inversion of red and green color features"

**Supplementary figures**

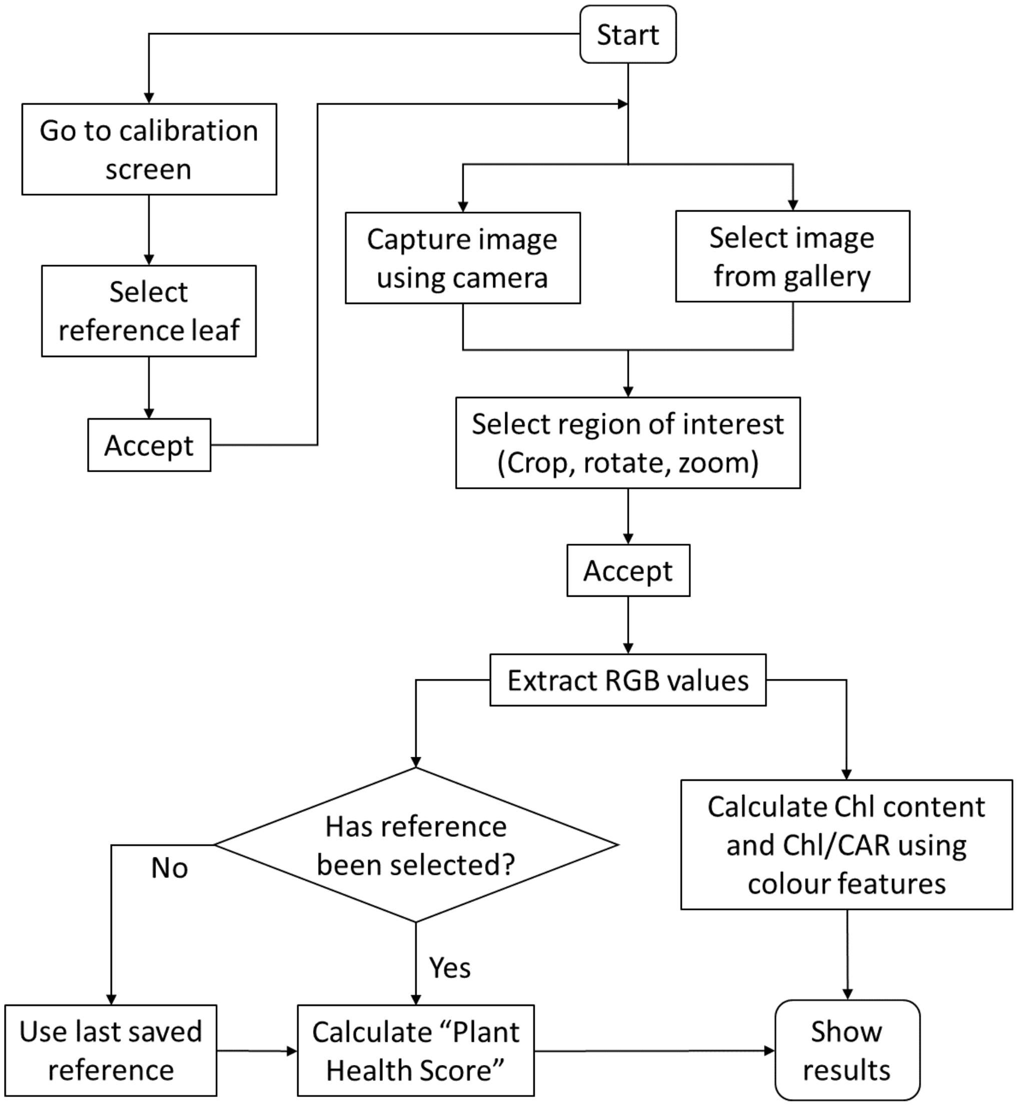


**Fig. S1** Algorithm of the "Plant Health Assessment" smartphone application.

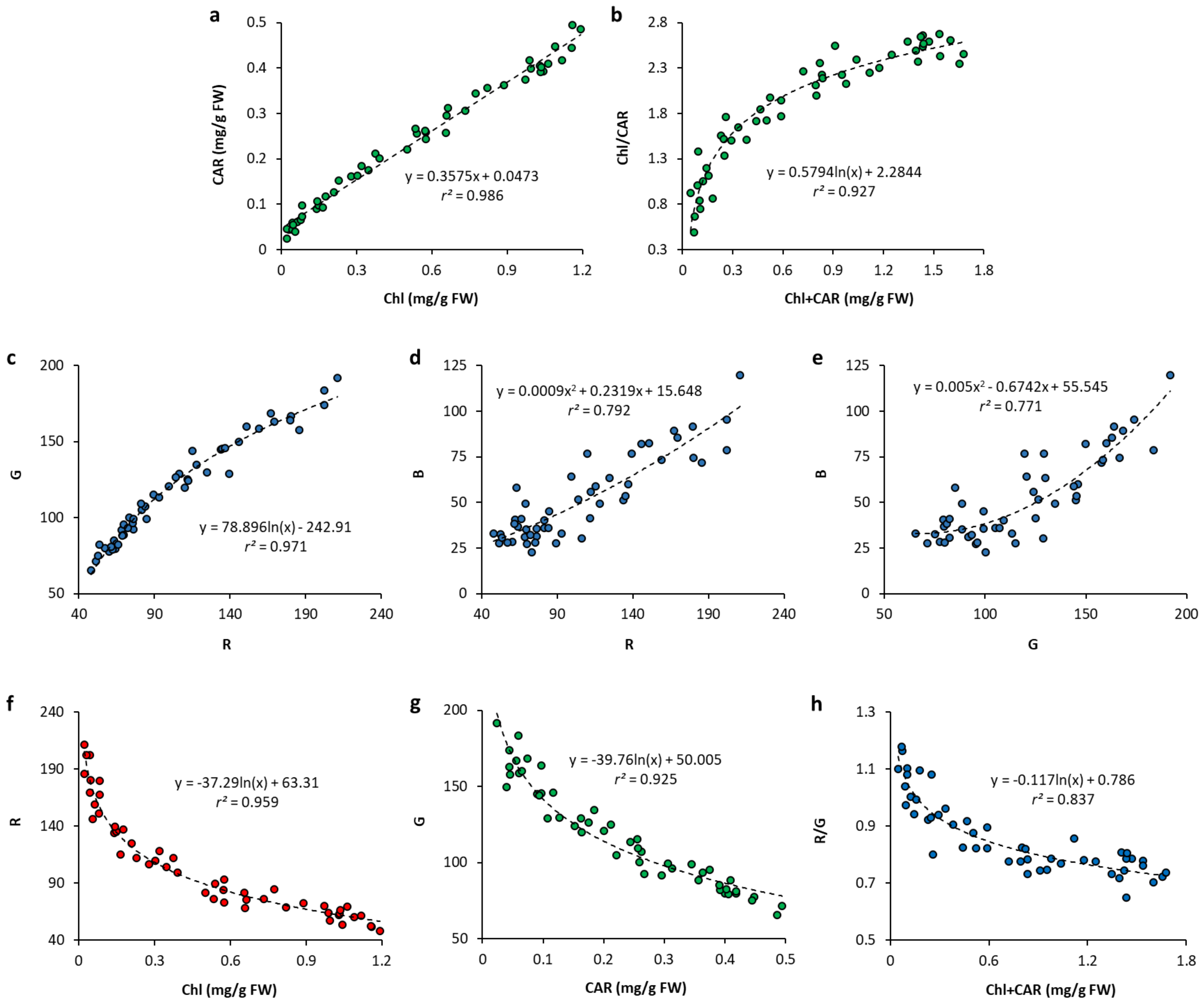


**Fig. S2** Relation between chlorophyll (Chl) and carotenoid (CAR) contents (a), Chl+CAR and Chl/CAR (b), red, green, and blue (RGB) color features of leaf digital images (c–e), Chl and R (f), CAR and G (g), and Chl+CAR and R/G (h) (*n* = 50, $p < 0.001$).

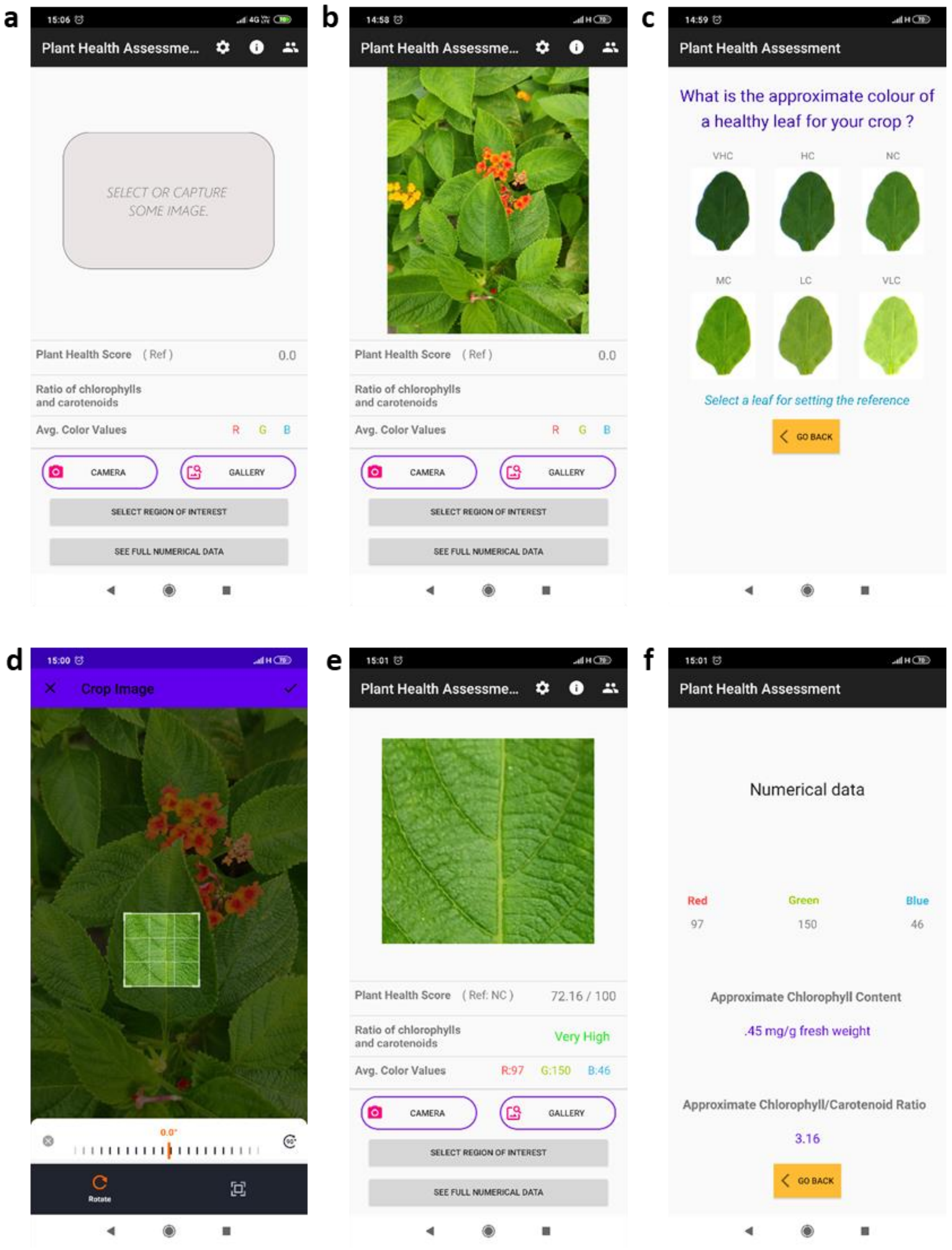


**Fig. S3** Screenshots of the "Plant Health Assessment" smartphone application user interface. The home screen (a), selected image (b), calibration screen (accessed via the cogwheel symbol) (c), selection of region of interest (d), home screen showing plant health score and RGB values (e), and the numerical data screen (f) have been shown in the order which must be followed for operating the application. In the calibration screen (c), the reference leaf with the colour of a healthy leaf of the target plant species must be selected to get the plant health score. The screen for selecting the region of interest provides the option for zooming, rotating, and cropping the image. VHC = Very high chlorophyll; HC = High chlorophyll; NC = Normal chlorophyll; MC = Medium chlorophyll; LC = Low chlorophyll; VLC = Very low chlorophyll.
